# Alcohol Disrupts Neural Differentiation Through Endoplasmic Reticulum Stress and PERK Pathway Activation

**DOI:** 10.1101/2025.05.16.654531

**Authors:** Zuohui Zhang, Wen Wen, Hong Lin, Di Hu, Hui Li, Jia Luo

**Author notes:** Correspondence author: Jia Luo, Department of Pathology, University of Iowa Carver College of Medicine, Iowa City, IA 52242, USA.

## Abstract

Prenatal alcohol exposure (PAE) can lead to fetal alcohol spectrum disorder (FASD), a condition marked by developmental brain defects that result in neurobehavioral and cognitive impairments. However, the underlying molecular mechanisms remain poorly understood. Brain development is a highly regulated process, with neurogenesis playing a crucial role. A key stage in this process is neural differentiation, which is essential for proper brain function. This study aims to investigate how alcohol disrupts neural differentiation. NE-4C cells, a neural stem cell line derived from the mouse embryonic brain, were utilized as an *in vitro* model. As an *in vivo* model, pregnant mice were exposed to alcohol between gestation days 14 and 16, after which newly formed neurons in the ventricular zone (VZ) were analyzed. To examine the role of endoplasmic reticulum (ER) stress, tunicamycin (TM), and MANF-deficient NE-4C cells were employed. Neural differentiation was assessed using immunofluorescence, immunoblotting and flow cytometry. Alcohol impaired the differentiation of NE-4C cells into neurons and astrocytes without impacting cell migration. It also induced ER stress, preferably activating the PERK pathway. Similarly, ER stress caused by TM and MANF deficiency disrupted neural differentiation and activated PERK. Inhibiting PERK mitigated alcohol-induced impairment of neuronal differentiation. PAE decreased the number of newly formed neurons in the VZ of fetal brain while having little effects on cell survival and proliferation. Inhibiting PERK partially reversed the reduction of new neurons caused by PAE. Thus, alcohol-induced ER stress, particularly PERK activation, may contribute to impaired neurogenesis linked to FASD.

## Introduction

In the U.S., 13.5% of adults report consuming alcohol during pregnancy, with 5.2% engaging in binge drinking within the past 30 days [1]. Prenatal alcohol exposure (PAE) is a major risk factor for fetal alcohol spectrum disorders (FASDs), which are characterized by abnormal brain development and impaired behavior. Neuroimaging studies of children and adolescents with FASDs reveal that PAE leads to significant changes in brain morphology, including reduced brain size and weight [2]. Alcohol can cross the placenta and directly affect the developing fetal brain, impairing neurogenesis—one of the key processes responsible for normal brain development. However, the cellular and molecular mechanisms underlying these effects remain poorly understood.

Neurogenesis is a complex and tightly regulated process within the developing central nervous system (CNS), involving neural stem cell (NSC) proliferation, migration, and differentiation. This process depends on efficient protein synthesis, which enables NSCs to transition from a quiescent to an active state. Therefore, maintaining protein homeostasis (proteostasis) is essential for neurogenesis. Proteostasis encompasses protein synthesis, folding, modification, degradation, and transport, with the endoplasmic reticulum (ER) serving as a critical hub for these functions. When proteostasis is disrupted and unfolded or misfolded proteins accumulate in the ER, ER stress occurs, triggering the unfolded protein response (UPR). The UPR primarily functions to restore balance by increasing ER chaperone expression, enhancing misfolded protein degradation, and temporarily reducing protein synthesis [3]. However, when ER stress becomes severe and homeostasis cannot be restored, the UPR shifts toward apoptotic cell death [3, 4]. The UPR is primarily regulated by three key ER transmembrane proteins: PKR-like ER kinase (PERK), inositol-requiring enzyme 1α (IRE1α), and activating transcription factor 6 (ATF6).

Growing evidence suggests that alcohol exposure disrupts proteostasis and induces ER stress in multiple organ systems, including the liver, pancreas, heart, and lungs [5–9]. Our previous studies, using both *in vitro* and *in vivo* models, demonstrated that alcohol exposure also induces ER stress in neurons within the developing and mature CNS [10–14]. However, whether alcohol-induced ER stress contributes to impaired neurogenesis in the developing brain remains unclear. In this study, we explored the role of ER stress in alcohol-mediated disruption of neural differentiation, using an *in vitro* model of cultured mouse NSCs and an *in vivo* model of the fetal mouse brain. Our findings reveal that alcohol exposure impairs neuronal differentiation through ER stress, specifically via activation of the PERK pathway. More importantly, both *in vitro* and *in vivo* experiments demonstrated that inhibiting the PERK pathway mitigated alcohol-induced impairment of neuronal differentiation.

## Materials and Methods

### Materials

MEM (11095-080), FBS, antibiotic-antimycotic (15240112), MTT (M5655), anhydrous DMSO (276855), all-trans RA (R2625), DAPI (D9542), Poly-L-lysine hydrobromide (P6282), and GSK2606414 (516535) were obtained from Sigma-Aldrich (St. Louis, MO, United States); PFA (15714) was obtained from Electron Microscopy Sciences (Hatfield, PA, United States); DC protein assay kit (5000112) was obtained from Bio-Rad Laboratories (Hercules, CA, United States); GeneArt Genomic Cleavage Detection Kit (A24372), and Lipofectamine 3000 Reagent (L3000008) were obtained from Life Technologies (Carlsbad, CA, United States; control CRISPR/Cas9 plasmid (sc-418922), mouse ARP double nickase plasmid (sc-428989-NIC), UltraCruz transfection reagent (sc-395739), and plasmid transfection medium (sc-108062) were obtained from Santa Cruz Biotechnology (Dallas, TX, United States); pGEM-T-easy vector (A1360) was obtained from Promega (Madison, WI, United States); 10-beta competent E. coli (C3019I) was obtained from New England Biolabs (Ipswich, MA, United States); VECTASHIELD mounting medium (H-1400 and H-1500) was obtained from Vector Laboratories (Burlingame, CA, United States).

Anti-β-III-tubulin (ab18207), anti-ARMET/ARP (MANF) (ab6727) and anti-phospho-IRE1(ab48187) antibodies were obtained from Abcam (Cambridge, MA, United States); Anti-ATF4 (CST11815), anti-phosphor-PERK (CST3179), anti-PERK (CST3192), anti-phosphor-eIF2a(CST3398), anti-eIF2a (CST9722), anti-IRE1(CST3294), anti-GFAP (CST3670), anti-SSEA-1 (CST4744), anti-DCX (CST4604), anti-Ki67 (CST 12202S) and anti-β-Actin (CST3700) antibodies were obtained from Cell Signaling Technology (Danvers, MA, United States); Anti-ATF6 (NBP1-40256) was obtained from Novus Biologicals Company (Littleton, CO, United States); Anti-GFAP antibody (13-0300) was obtained from Invitrogen (Waltham, MA, United States); Alexa Fluor® 488 anti-β-3-Tubulin (TUBB3) (657404), Alexa Fluor® 488 Mouse IgG2a, κ Isotype Ctrl (ICFC) (400238), APC anti-mouse/human CD15 (SSEA-1) (125618), and APC Mouse IgM, κ Isotype Ctrl (401616) antibodies were obtained from Biolegend (San Diego, CA, United States); Secondary antibodies conjugated to horseradish peroxidase (NA931V and NA934V) were obtained from GE Healthcare Life Sciences (Piscataway, NJ, United States); Alexa-488 conjugated anti-mouse (A21202), Alexa-594 conjugated anti-mouse (A11005), Alexa-488 conjugated anti-rabbit (A21206) and Alexa-594 conjugated anti-rabbit (A11012) antibodies were obtained from Life Technologies (Carlsbad, CA, United States).

## Methods

### Animal model and alcohol exposure

All experimental procedures were approved by the Institutional Animal Care and Use Committee (IACUC) at the University of Iowa and performed following regulations for the Care and Use of Laboratory Animals set forth by the National Institutes of Health (NIH) Guide. To investigate of the effects of alcohol on brain development, pregnant C57BL6 mice (The Jackson Laboratory, Bar Harbor, ME) were randomly assigned to two groups: control and alcohol group. They were administered water or alcohol (EtOH: 5 g/kg) by oral gavage once per day on gestation days (GD) 14 and 16. The mice were sacrificed, and fetal brains were collected on GD17. For inhibition of PERK pathway with GSK2606414, pregnant mice were randomly assigned to four groups: control, GSK2606414, alcohol and alcohol plus GSK2606414 group. The mice were pretreated with GSK2606414 (50 mg/kg) or vehicle 1 hour before the administration of water or alcohol (5 g/kg) by oral gavage once per day on GD14 and 16. The mice were sacrificed, and fetal brains were collected on GD17.

### Cell culture, neural differentiation and measurement of neurospheres

NE-4C (CRL-2925) cells were from ATCC (Manassas, VA, United States). NE-4C cells were maintained in MEM supplemented with 10% FBS and 1% antibiotic-antimycotic at 37°C in a humidified incubator with 5% CO₂. To induce neural differentiation, cells were seeded in 6-well plates at a density of 2 × 10L cells per well and cultured for 24 hours. The growth medium was then carefully replaced with differentiation medium consisting of MEM supplemented with 5% FBS and 1.5 × 10⁻L M all-trans RA for 2 days. Following this, cells were maintained in RA-free MEM with 10% FBS until further analysis. For alcohol treatment, NE-4C cells were cultured in complete MEM medium containing 0.4% (w/v) ethanol. To induce ER stress, NE-4C cells were incubated with 30 ng/ml tunicamycin for the time periods indicated in each figure. To determine the appropriate concentration of GSK2606414, the compound was added to the culture medium at concentrations of 0.1, 1, 10, 50, and 100 µM, 30 min prior to alcohol and tunicamycin treatment. To quantify the size of neutrospheres, at least five randomly selected fields per well were imaged using an Olympus BX51 light microscope with a 40X objective. The five fields were chosen in an X pattern, with four at the corners and one in the center of the well. The size of neurospheres was measured using ImageJ software.

### CRISPR/Cas9 knockout of mesencephalic astrocyte-derived neurotrophic factor (MANF)

The knockout of MANF in NE-4C cells was carried out using the CRISPR/Cas9 technique according to a previously described method [19]. To establish knockout single-cell colonies, we utilized two CRISPR/Cas9 plasmids: a control plasmid (referred to as’control CRISPR’) and a targeted mouse MANF double-nickase plasmid (referred to as’MANF CRISPR’). The control CRISPR plasmid carried a GFP marker for transfection verification, while MANF CRISPR consisted of two plasmids targeting exon 2 of the MANF gene—one encoding a puromycin resistance gene and the other containing a GFP marker. 4×10^5^ NE-4C cells were seeded in 6-well plates and cultured to approximately 70% confluence before transfection with 1 µg of either control CRISPR or MANF CRISPR. Transfection was performed using the UltraCruz transfection reagent and plasmid transfection medium, following the manufacturer’s guidelines. Forty-eight hours after transfection, GFP-positive cells were sorted using flow cytometry. To enhance selection, cells transfected with MANF CRISPR were further treated with 1 µg/mL puromycin for five days before a second round of flow cytometry sorting to isolate single cell colonies. To validate successful biallelic MANF knockout, protein lysates from each colony were subjected to immunoblotting (IB) analysis of MANF expression.

### Wound healing assay

NE-4C cells were seeded in 6-well plates at a density of 4 × 10L cells per well. After 24 hours, a linear wound was created by scratching the cell monolayer with a sterile pipette tip. The wells were gently washed with PBS to remove detached cells. Phase-contrast images of the wound area were captured at 0, 24, 48, and 72 hours after scratching to monitor cell migration and wound closure.

### Protein extraction and IB

NE-4C cells or fetal brain tissues were washed with ice-cold PBS and lysed on ice for 15 minutes in RIPA buffer containing 150 mM NaCl, 1 mM EGTA, 50 mM Tris– HCl (pH 7.5), 0.5% Nonidet P-40 (NP-40), and 0.25% SDS, with freshly added protease inhibitors: 5 mg/mL leupeptin, 5 mg/mL aprotinin, 3 mM sodium orthovanadate, and 0.3 mg/mL PMSF. The lysates were centrifuged at 10,000 × g for 30 minutes at 4°C, and the resulting supernatant was collected. Protein concentrations were measured using the DC protein assay (Bio-Rad Laboratories) following the manufacturer’s instructions. Approximately 20–30 μg of total protein from each sample was separated by SDS-PAGE. The resolved proteins were transferred onto nitrocellulose membranes and incubated overnight at 4°C with specific primary antibodies. The primary antibodies and their final dilutions were as follows: anti-ARMET/ARP (MANF) (1:1000), anti-phospho-PERK (1:1000), anti-PERK (1:1000), anti-ATF4 (1:1000), anti-ATF6 (1:1000), anti-phospho-eIF2α (1:1000), anti-eIF2α (1:1000), anti-phospho-IRE1 (1:1000), anti-IRE1 (1:1000), anti-β-III-tubulin (1:1000), anti-GFAP (1:1000), anti-MANF (1:1000), and anti-β-actin (1:10000). After washing with TBST, membranes were incubated with horseradish peroxidase-conjugated secondary antibodies (1:5000) for 1 hour at room temperature. Protein bands were visualized using the Amersham ECL Prime Western Blotting Detection Reagent (GE Healthcare Life Sciences), and band intensities were quantified using Image Lab software (Bio-Rad Laboratories).

### Immunofluorescence (IF)

NE-4C cells were plated onto 24-well plates containing sterile coverslips pre-coated with 15 μg/mL poly-L-lysine. Following differentiation, cells were fixed with 4% PFA and permeabilized using 0.25% Triton X-100 in PBS for 10 minutes at room temperature. Subsequently, cells were blocked with 1% BSA and 2% goat serum in PBS for 30 minutes before incubation with primary antibodies: anti-β-III-tubulin (1:100), anti-GFAP (1:200), for 1 hour at room temperature. After washes with PBS, cells were incubated with Alexa Fluor-conjugated secondary antibodies (1:200). Cell nuclei were counterstained with DAPI. Fetal mouse brains were fixed with 4% PFA, cryoprotected in 30% sucrose, and processed for IF staining as previously described [10]. The brain sections were blocked with 1% BSA at room temperature. Doublecortin (DCX) in the ventricular zone (VZ) of mouse brain was immunostained with anti-DCX antibody (1:1600); Ki-67 in the VZ of mouse brain was immunostained with anti-Ki-67 antibody (1:500) at 4 °C overnight. The sections were then incubated with secondary antibody conjugated to Alexa Fluor 488 or 594. Cell nuclei were counterstained with DAPI. Fluorescence images were acquired using an Olympus IX81 inverted fluorescence microscope.

### Flow cytometry

Flow cytometry was used to quantitatively assess the percentage of NE-4C cells undergoing neuronal differentiation. The stage-specific embryonic antigen-1 (SSEA-1), also known as CD15, is a marker for NSCs [20, 21]. It is involved in the maintenance of stem cells, and its expression is downregulated upon neural stem cell differentiation. The cells were harvested and prepared as a single-cell suspension. They were washed twice with washing buffer (PBS containing 1% FBS and 0.1% NaN₃). The cells were then stained with APC-conjugated anti-mouse/human CD15 (SSEA-1) for 20–30 minutes on ice in the dark. After staining, the cells were washed with washing buffer and fixed with fixation buffer (4% PFA in washing buffer) for 20 minutes at room temperature in the dark. Following two washes with washing buffer, the cells were permeabilized with Intracellular Staining Perm Wash Buffer (00-8333-56, Invitrogen) for 20 minutes at room temperature. The cells were then resuspended and incubated with Alexa Fluor® 488-conjugated anti-β-III-Tubulin antibody (657404, Biolegend) for 30–60 minutes at room temperature. After two washes, the cells were resuspended in 0.5–1 mL of washing buffer. Fluorescence intensity was measured using a FACS (BD LSR II), and data acquisition and analysis were performed using FlowJo software.

### MTT Assay

Cell viability was determined by measuring cell metabolic activity using the MTT assay. NE-4C cells were seeded in 96-well plates at a density of 2 × 10³ cells per well. At the indicated time points, MTT was added to each well at a final concentration of 500 μg/mL and incubated at 37°C for 2 hours. Following incubation, the medium was carefully removed, and 100 μL of DMSO was added to dissolve the MTT formazan. Absorbance was measured at 595 nm using a Beckman Coulter DTX 880 Multimode Detector plate reader.

### Statistical analysis

Statistical analyses were performed using GraphPad Prism version 10. All data were presented as mean ± SEM from at least three independent experiments per group. Differences between experimental groups were evaluated using a student t-test, one-way or two-way ANOVA. A p value < 0.05 was considered as significant difference. Tukey’s post hoc test was applied following one-way ANOVA, while Bonferroni’s post hoc test was used following two-way ANOVA to analyze the difference among specific treatment groups.

## Results

### Alcohol impairs neural differentiation

NE-4C cells are a neural stem cell line derived from mouse embryonic brain vesicles [15]. They can differentiate to neuron and astrocyte when exposed to retinoic acid (RA) and are well-established model for studying neurogenesis [16–18]. First, we examined the effects of alcohol on NE-4C cell viability. NE-4C cells were treated with alcohol (0.2%-0.8%) for 24 hours or 72 hours. The MTT assay was performed to evaluate the cell viability (Fig 1). Alcohol exposure at 0.8% for 72 hours reduced the cell viability of NE-4C cells, while alcohol at 0.2% and 0.4% for 72 hours had little effect on cell viability (Fig.1B). As a result, alcohol concentration of 0.4% was used for the subsequent experiments. To investigate the effects of alcohol on neural differentiation, NE-4C cells were treated with RA in the presence or absence of alcohol. The expression of β-III-tubulin, a neuron marker was used for detection of neuronal differentiation of NE-4C cells [22, 23]. Upon RA treatment, NE-4C cells differentiated into neuronal cells as evident by the increased expression of β-III-tubulin (Fig. 1C). Alcohol exposure significantly decreased the expression of β-III-tubulin (Figs.1D, F-I). In addition, the length of neurites in alcohol-treated group was significantly shorter than that of controls (Fig.1E). The results indicated that alcohol impaired neuronal differentiation of NE-C cells.

**Figure 1.**
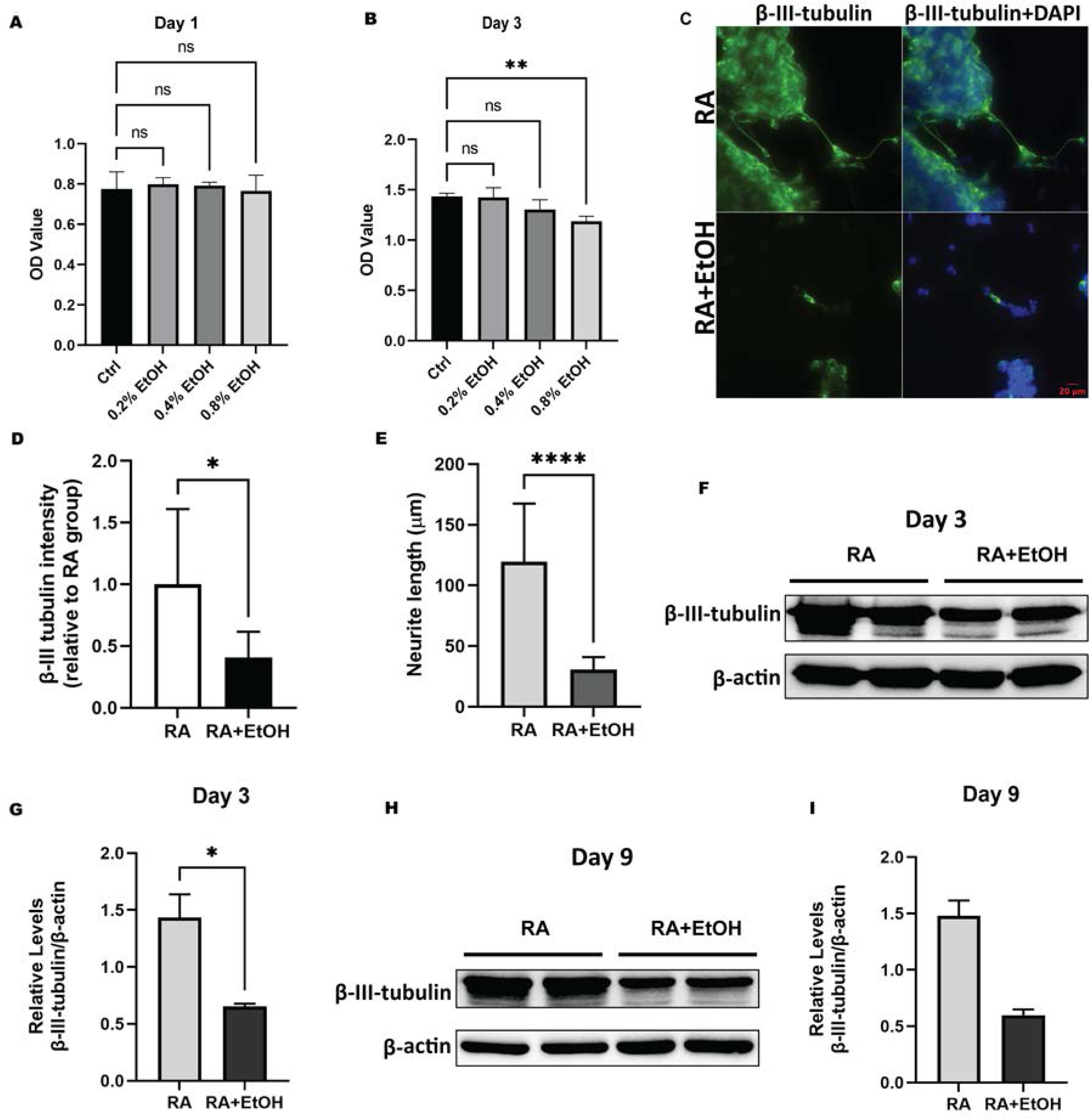
Effects of alcohol on neuronal differentiation. **A** and **B**: NE-4C cells were treated with different concentrations of alcohol (EtOH: 0.2%-0.8%). Cell viability was determined by MTT assay after 1 day (**A**) and 3 days of treatment (**B**). **C** and **D**: NE-4C cells were treated with RA (1.5 × 10⁻L M) in the presence or absence of 0.4% alcohol (EtOH). β-III-tubulin expression was detected by immunofluorescence (IF) after 3 days of treatment (**C**). The intensity of β-III-tubulin expression was determined (**D**). **E**: The length of neurites (**F**) was quantified as described in the Materials and Methods. **F-I**: The expression β-III-tubulin was also examined by immunoblotting (IB) analysis and quantified after 3 days (**F** and **G**) and 9 days of treatment (**H** and **I**). The relative protein levels were normalized to β-actin. Data are presented as mean ± SEM of three experiments (n=3). * *p* < 0.05, or ** *p* <0.01, or **** *p* < 0.0001 when compared to respective controls.

NE-4C cells treated with RA also generate astrocytes in later stages of differentiation [16]. To evaluate the effects of alcohol on NE-4C differentiation to astrocytes, we examined the expression of GFAP, a marker of astrocytes 9 days after the treatment of RA and alcohol. Alcohol significantly reduced the expression of GFAP, indicative of inhibition of glial differentiation (Fig. 2).

**Figure 2.**
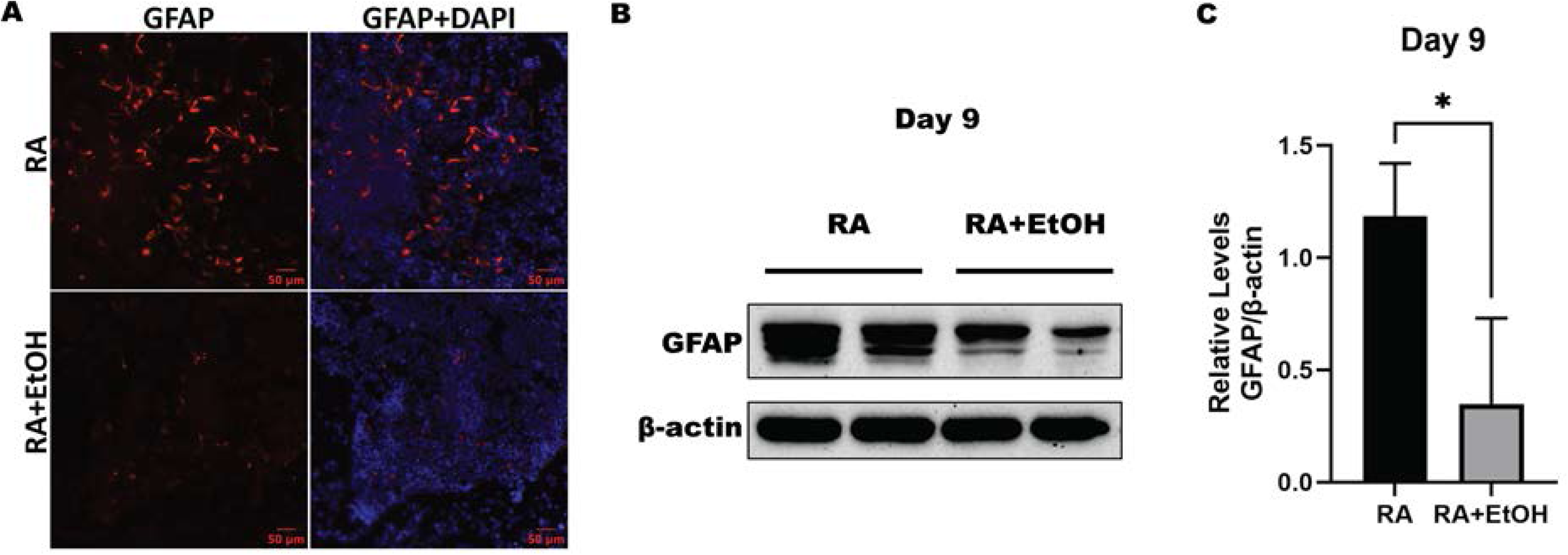
Effects of alcohol on astrocyte differentiation. **A** and **B**: NE-4C cells were treated with RA (1.5 × 10⁻L M) in the presence or absence of 0.4% alcohol (EtOH). The expression of GFAP examined by IF (**A**) and IB (**B**) after 9 days of treatment. **C**: The expression levels of GFAP levels were quantified and normalized with β-actin. Data are presented as mean ± SEM of three experiments. * *p* < 0.05?

### Alcohol does not impair migration of neural progenitor cells

Generation of NSCs as well as subsequent cell migration are important processes of neurogenesis crucial for proper brain development and function. Appropriate neuronal migration ensures that newly formed neurons are correctly positioned in the brain to establish functional neural circuits which is essential for brain development and function. We employed wound healing assay to investigate the impact of alcohol on cell migration. As shown in Fig. 3, alcohol had a minimal effect on the migration of NE-4C cells.

**Figure 3.**
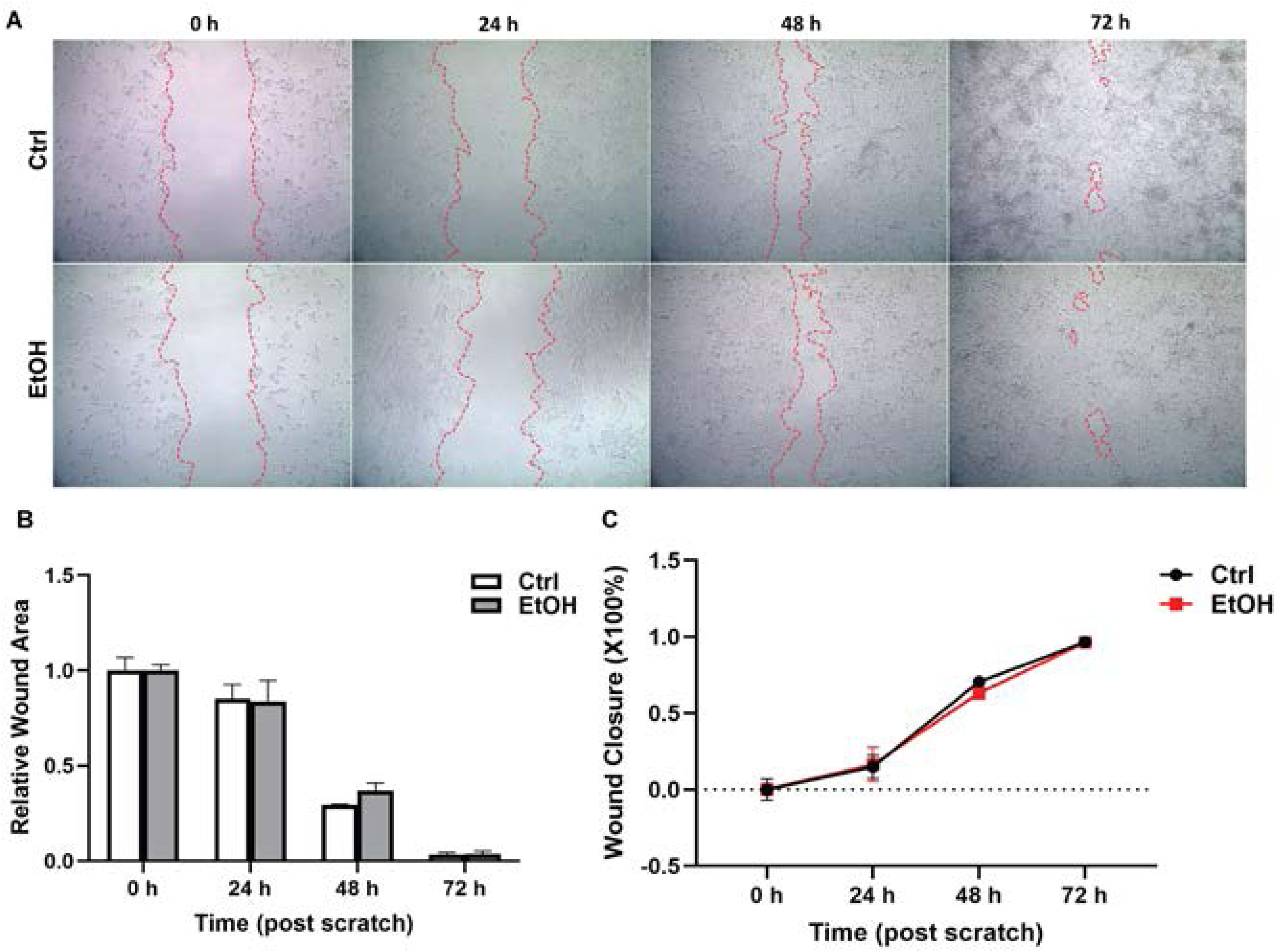
Effects of alcohol on cell migration. **A**: NE-4C cells were treated with or without 0.4% alcohol (EtOH). Representative images show cell migration in the wound healing assay at different time points after creating a scratch in the cell monolayer. **B**: The wound area relative to the initial area at 0 h was quantified. **C**: The wound closure which was expressed as the percentage of recovery of the wound area was quantified. Data are presented as mean ± SEM of three experiments.

### Alcohol induces ER stress in NSCs

To evaluate the role of ER stress in alcohol-induced impairment of neural differentiation, we first determined whether alcohol induced ER stress in NE-4C cells and examined the activation of UPR pathways. Alcohol activated PERK pathway which was evident by the increased phosphorylation of PERK and eIF2a, and the upregulation of AFF4 (Fig. 4). In addition, alcohol also stimulated IRE1 and ATF6 pathways and increased the expression of MANF, but to a lesser extent (Fig. 4). The results indicated that alcohol induced ER stress in NE-4C cells.

**Figure 4.**
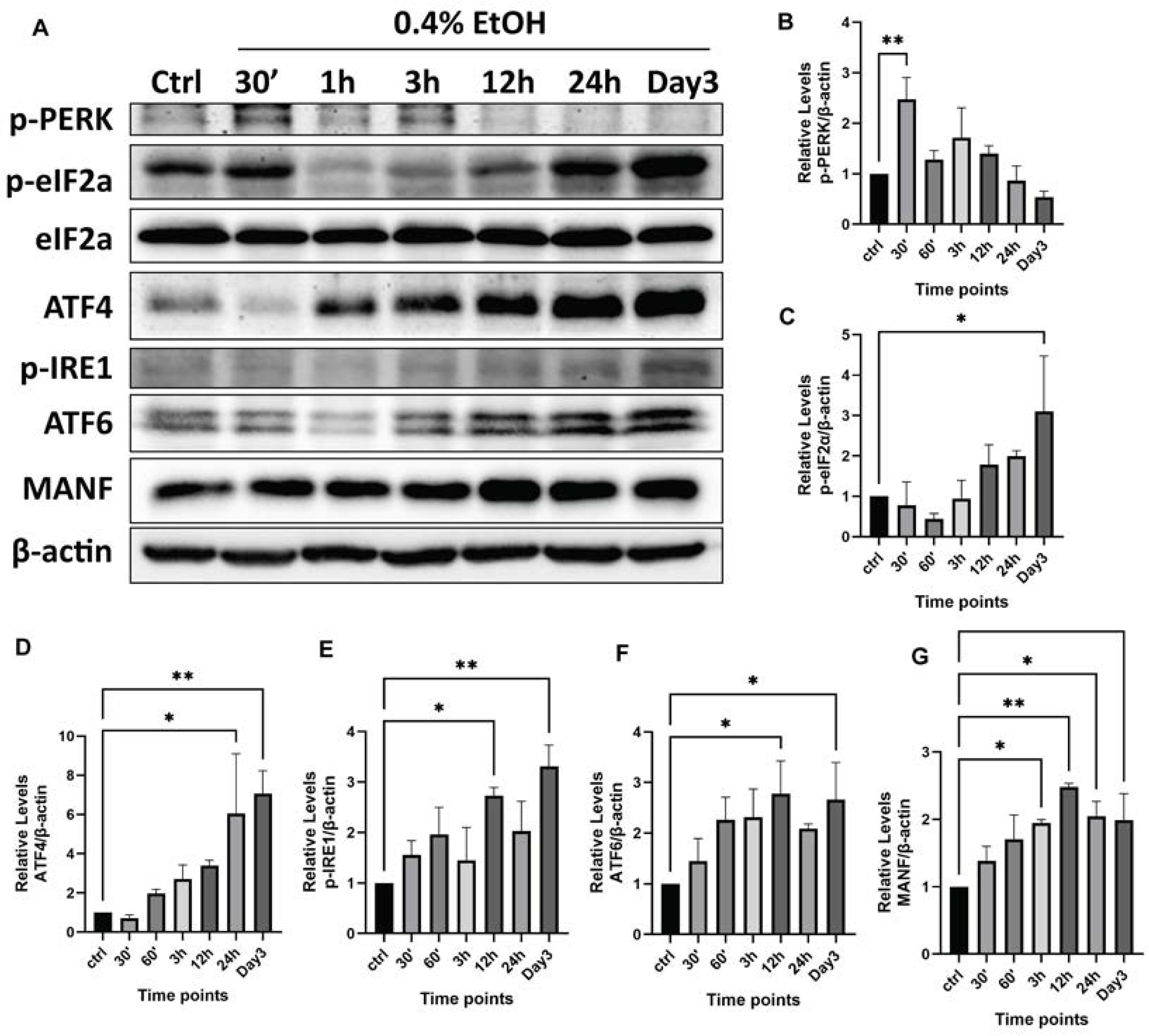
Effects of alcohol on ER stress. **A**: NE-4C cells were treated with 0.4% alcohol for 0.5, 1, 3, 12, 24, and 72 hours. Protein was extracted and subjected to IB analysis of phosphorylated PERK (p-PERK), p-eIF2α, eIF2α, ATF4, p-IRE1, ATF6, and MANF. **B-F**: The expression of these UPR proteins was quantified and normalized with β-actin. The experiments were replicated 3 time (n = 3). Data are presented as mean ± SEM. * *p* < 0.05, or ** *p* <0.01, when compared to controls (Ctrl).

### Tunicamycin (TM) impairs neuronal and glial differentiation

To determine the role of ER stress in neural differentiation, we employed a well-defined ER stress inducer, TM which is commonly used for studying ER stress-associated cellular alterations. TM inhibits the first step of N-linked glycosylation, leading to the accumulation of misfolded proteins, which triggers ER stress. We first determined the effect of TM on cell viability using a range of TM concentrations (Supplementary Fig. 1). Since TM of 30 ng/ml did not significantly affect the viability of NE-4C cells, we used this concentration in the subsequent experiments. We showed that TM induced ER stress as evident by the activation of three UPR pathways—PERK/ eIF2a, IRE1, and ATF6 as well as upregulation of MANF (Fig. 5). It appeared that TM affected PERK/eIF2a/ATF4 pathway more than other pathways in NE-4C cells. TM treatment significantly decreased the expression of β-III-tubulin (Fig.6A, and B, E-I). In addition, the length of neurites in TM-treated group was significantly shorter than that of controls (Fig.6C and D). The results indicated that TM impaired neuronal differentiation of NE-4C cells. TM also significantly reduced the expression of GFAP, indicative of inhibition of glial differentiation (Fig. 6G, I and J). Thus, like alcohol, TM inhibited neural differentiation.

**Figure 5.**
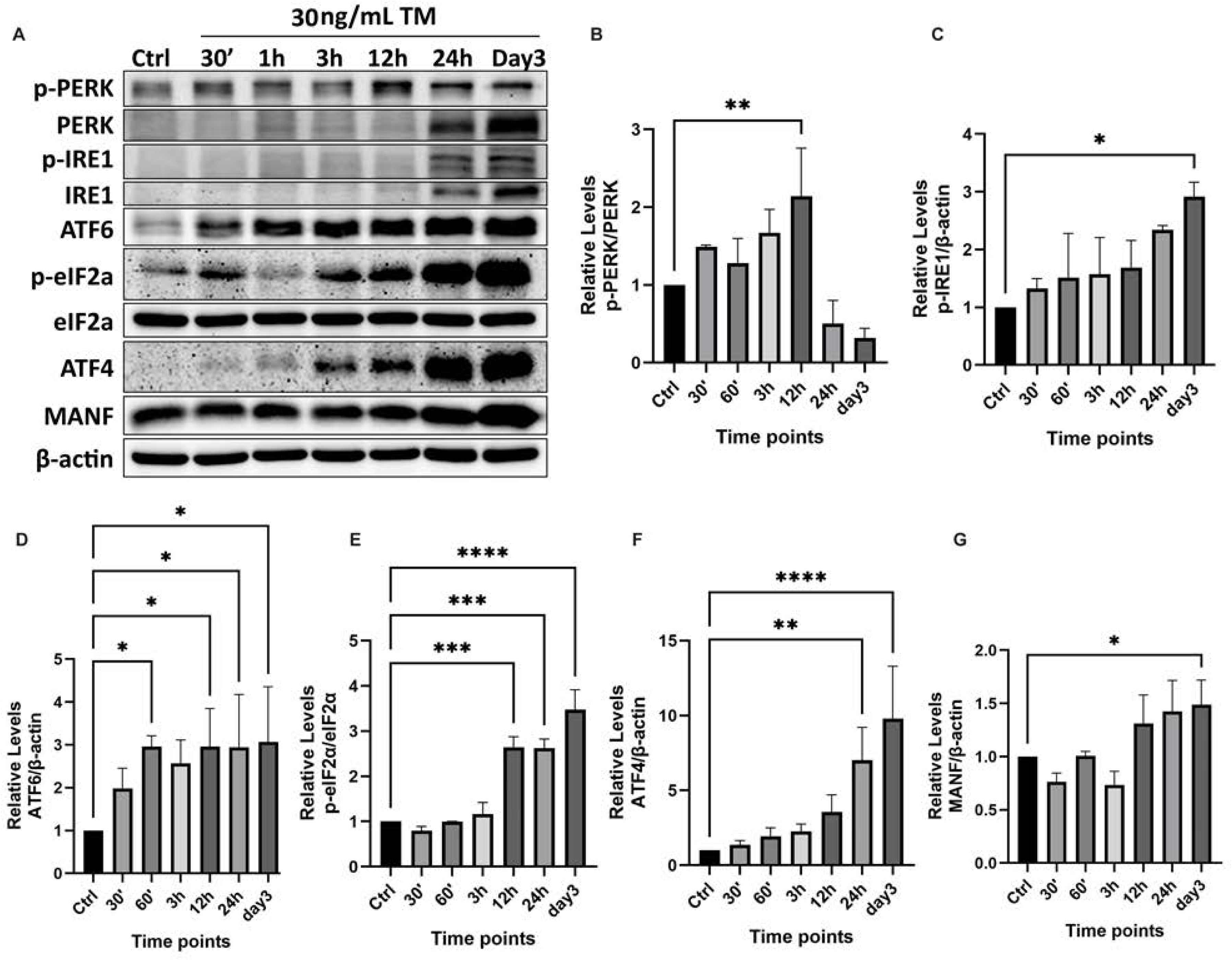
Effects of tunicamycin (TM) on ER stress. NE-4C cells were treated with TM (30 ng/mL) for indicated times, the expression of UPR proteins was analyzed by IB (**A**) and quantified (**B-G**) as described in Figure 4. The experiments were replicated three times. Data are presented as mean ± SEM. *p < 0.05, **p < 0.01. ***p < 0.001, ****p < 0.0001 when compared to controls.

**Figure 6.**
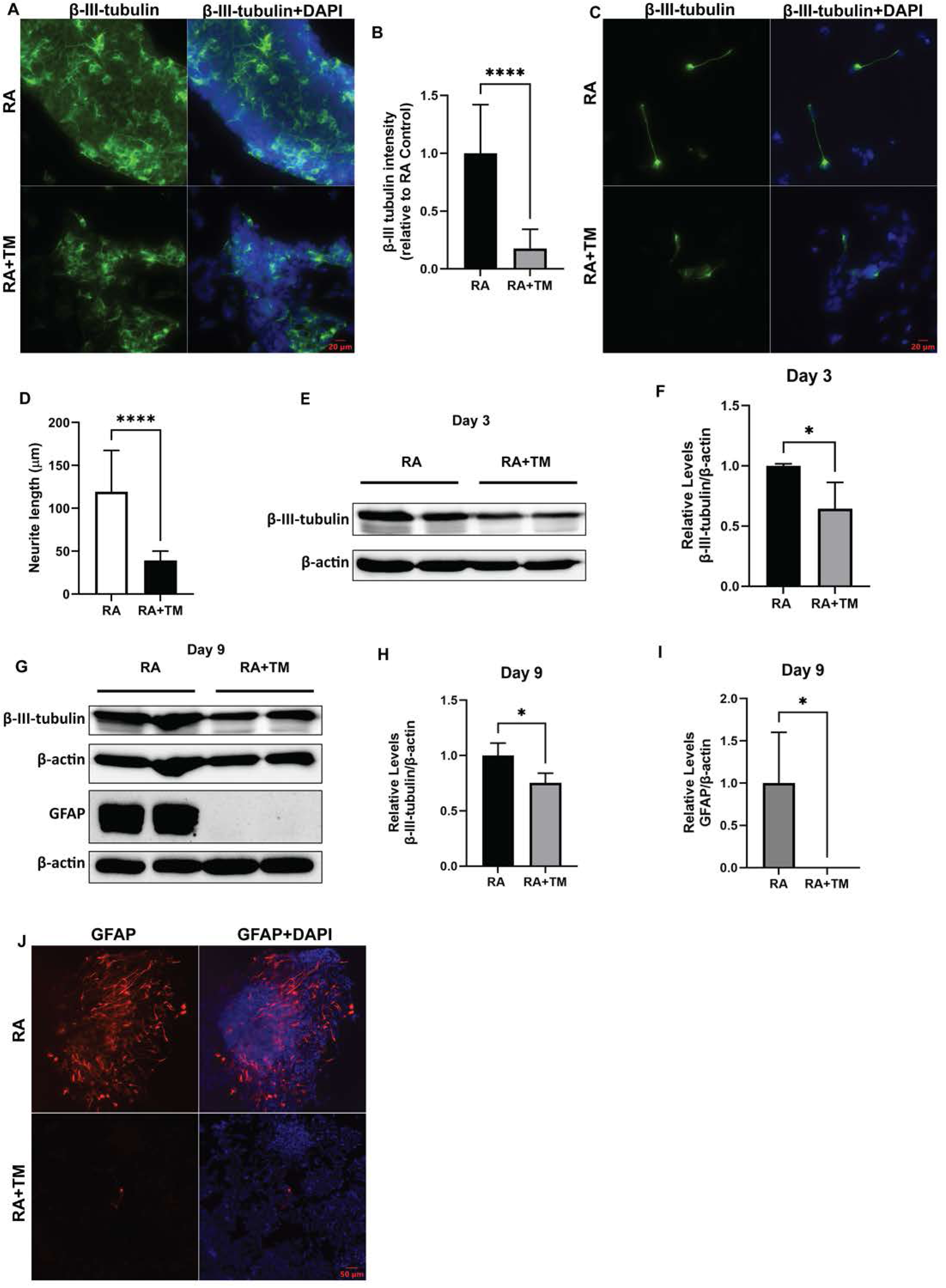
Effects of TM on RA-induced neuronal and astrocyte differentiation. **A**: NE-4C cells were treated with RA (1.5 × 10⁻L M) in the presence or absence of TM (30 ng/mL) for 48 hours. The expression of β-III-tubulin in the cultures was analyzed with IF on day 3. **B**: The intensity of β-III-tubulin expression was quantified as described in Materials and Methods. **C** and **D**: The neurites were visualized by IF and the length of neurites was determined. **E**: The expression of β-III-tubulin was determined by IB on day 3 after RA treatment. **F**: The expression of β-III-tubulin was quantified and normalized to β-actin. **G**: The expression of β-III-tubulin and GFAP was determined by IB on day 9 after RA treatment. **H** and **I**: The expression of β-III-tubulin and GFAP shown in panel GF was quantified and normalized to β-actin. **J**: The expression of GFAP in the cultures was analyzed with IF on day 9 after RA treatment. Data are presented as mean ± SEM of three replications. *p < 0.05, **** p< 0.0001, when compared to controls.

### MANF Deficiency induces ER stress and impairs neuronal and glial differentiation

MANF is an ER-resided protein that regulates ER homeostasis. MANF is upregulated under ER stress and functions to alleviate ER stress by interacting with components of the UPR [24, 25]. Increasing evidence indicates that MANF is critically involved in many ER stress-related diseases with protective effects [26–28]. Our previous research and that of others indicates that MANF knockout (KO) activates ER stress in neurons [12]. As an additional approach to investigate the role of ER stress in neural differentiation, we used CRISPR/Cas9 technique to knock out MANF in NE-4C cells. Two clones of MANF KO NE-4C cells were selected for the subsequent experiments (Fig. 7A). MANF KO in NE-4C cells induced ER stress and drastic activation of PERK and eIF2a pathway (Figs. 7B-E). The neuronal differentiation of MANF KO NE-4C cells was assessed by the expression of β-III-tubulin by IF staining, IB and flow cytometry. MANF KO significantly reduced the expression of β-III-tubulin in response to RA treatment (Figs. 7F-J). Furthermore, the length of neurites in MANF KO cells was much shorter than that of controls (Fig. 7K). Taken together, these findings indicated that ER stress induced by MANF deficiency impaired neuronal differentiation. Next, we examined the effect of MANF KO on glial differentiation on day 9 after RA treatment. The expression of GFAP was significantly lower in MANF KO NE-4C cells than that of controls (Fig. 8), suggesting that ER stress induced by loss of MANF also disrupted glial differentiation.

**Figure 7.**
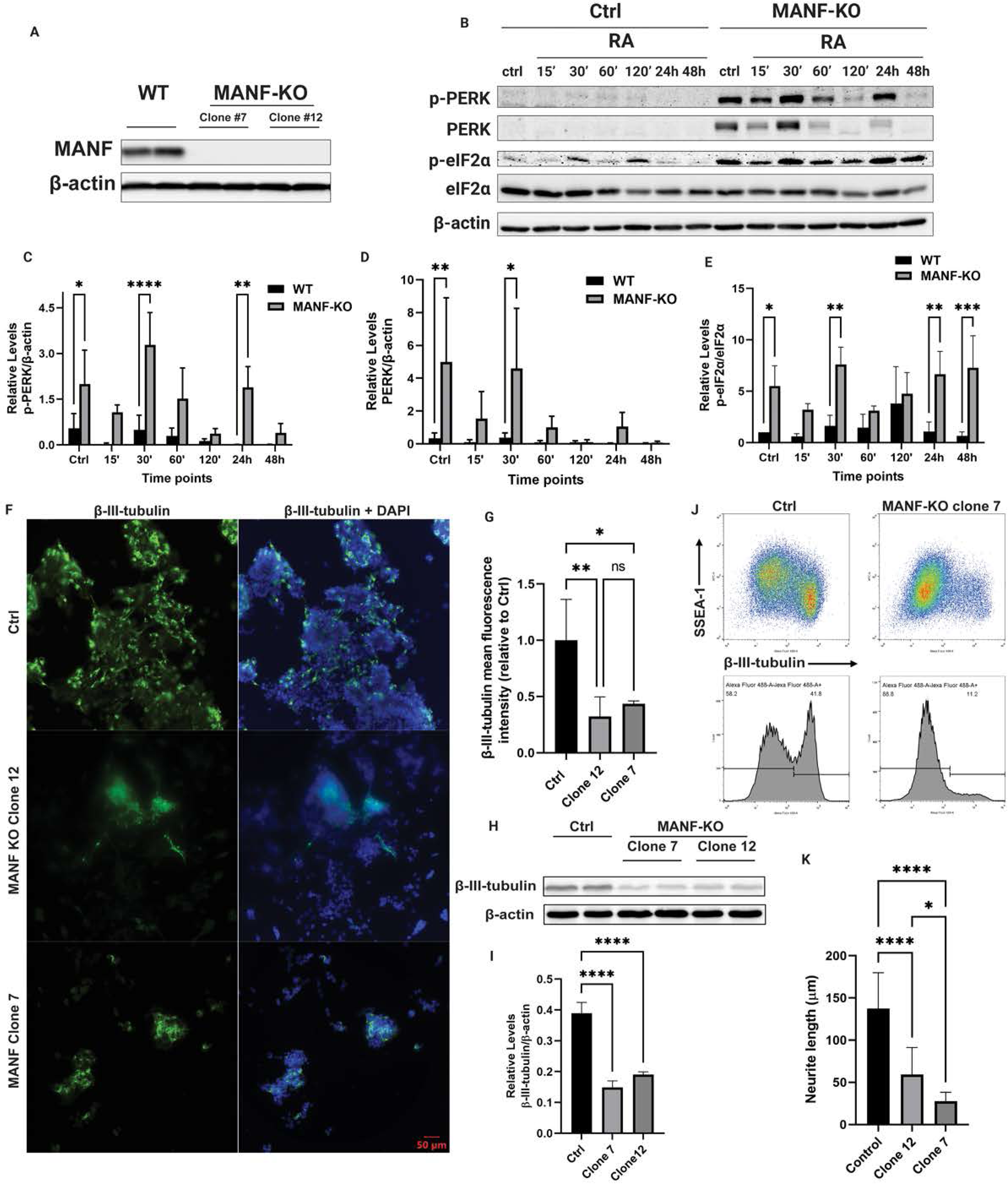
Effects of MANF Deficiency on ER stress and neuronal differentiation. **A**: MANF was knocked out (KO) in NE-4C cells as described in Materials and Methods. MANF KO was verified by IB and the results of two clones (Clone #7 and 12) were presented. **B**: Control and MANF KO NE-4C cells were treated with RA (1.5 × 10⁻L M) for 15’, 30’, 60’, 120’, 24 h, and 48 h. After that, protein was extracted and subjected to IB analysis of p-PERK, PERK, p-eIF2α, and eIF2α. **C-E**: The expression levels of these UPR proteins were quantified and normalized with β-actin. **F**: Control and MANF KO cells were treated with RA (1.5 × 10⁻L M) for 48 hours. The expression of β-III-tubulin was visualized by IF on day 3. **G**: The fluorescence intensity was quantified. **H**: The expression of β-III-tubulin was analyzed by IB after treatment of RA for 48 hours. **I**: The expression of β-III-tubulin was quantified and normalized to β-actin. **J**: The expression of SSEA-1 and β-III-tubulin was analyzed with flow cytometry as described in Materials and Methods. **K**: The length of neurites was quantified. Data are presented as mean ± SEM of three independent experiments. *p < 0.05, **p < 0.01, ***p < 0.001, ****p < 0.0001 when compared to controls.

**Figure 8.**
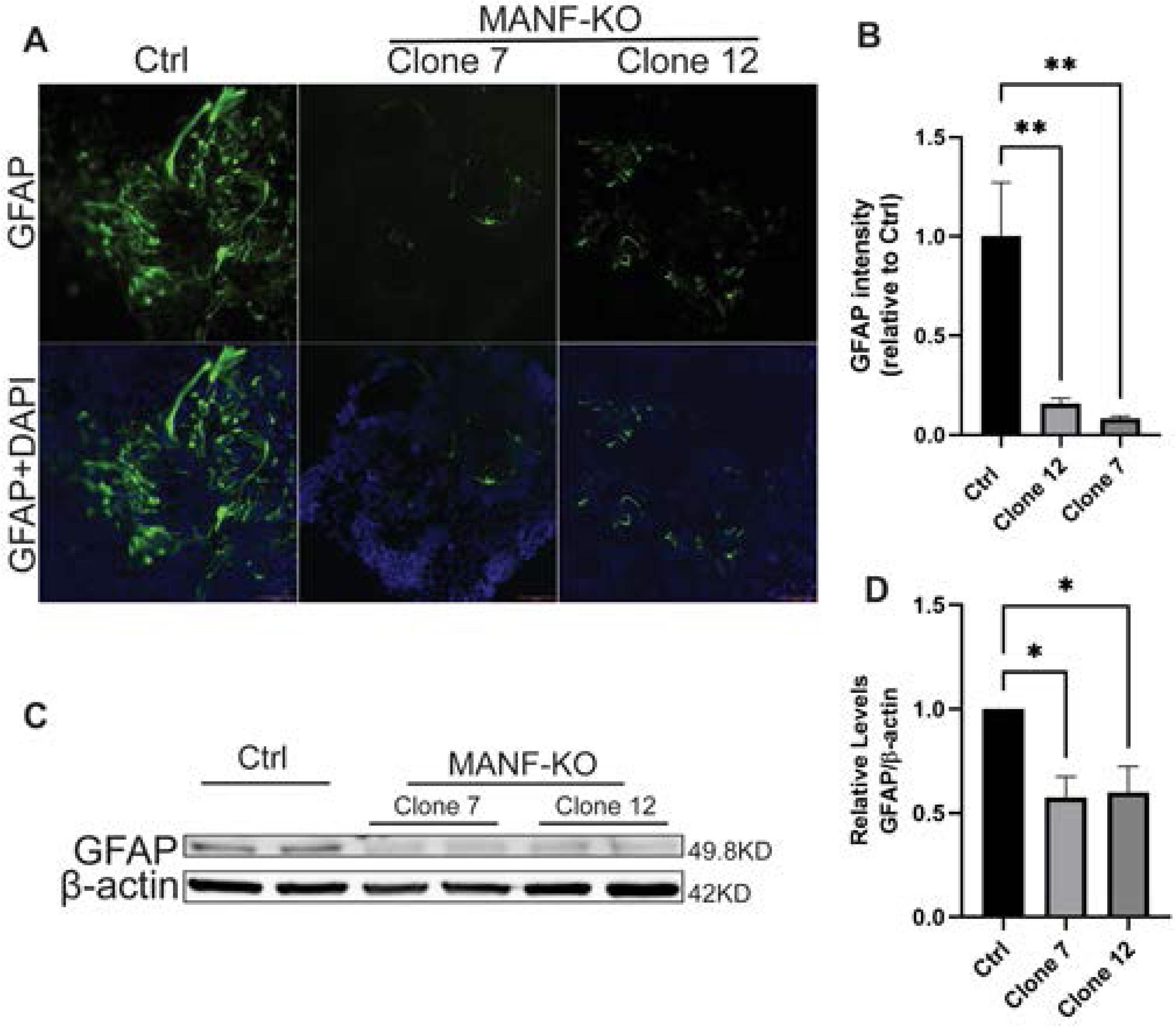
Effects of MANF Deficiency on glial differentiation. **A**: Control and MANF KO NE-4C cells (Clone 7 and 12) were treated with RA (1.5 × 10⁻L M) for 48 hours. The expression of GFAP in the cultures was visualized by IF on day 9. **B**: The intensity of GFAP signals in panel A was quantified. **C**: The expression of GFAP was determined by IB. **D**: The expression of GFAP was quantified and normalized to β-actin. Data are presented as mean ± SEM three independent experiments. *p < 0.05, **p < 0.01 when compared to controls.

### Inhibiting PERK mitigates neuronal differentiation impairments caused by TM and alcohol

The above results support the notion that alcohol activated ER stress which may impair neural differentiation. It appeared that PERK/eIF2a was preferably affected in NE-4C cells under ER stress condition. We therefore sought to determine whether PERK played a role in neural differentiation. We inhibited PERK activity using a specific PERK inhibitor GSK2606414 in NE-4C cells. GSK2606414 has been commonly used *in vitro* and *in vivo* to effectively inhibits PERK activity [29, 30]. We examined a range of GSK2606414 concentrations and determined that 1 μM had minimal toxicity but effectively inhibited PERK activity (Supplementary Fig. 2 and Fig. 9A and B). First, we determined the effects of GSK2606414 on TM-induced impairment of neural differentiation. The formation of neurospheres has been used to determine neuronal differentiation of NSCs [31–33]. TM significantly inhibited the formation of neurospheres (Fig. 9C and D). However, in GSK2606414-and TM-treated group, the formation neurospheres was significantly larger than TM-treated group (Fig. 9C and D). In addition, the length of neurites in GSK2606414-and TM-treated group was significantly longer than that of TM-treated group (Fig. 9G). GSK2606414 also mitigated TM-induced reduction of β-III-tubulin expression (Fig. 9E and F). The results were further confirmed by flow cytometry, which demonstrated an increased percentage of β-III-tubulin positive cells in in GSK2606414-and TM-treated group compared to TM-treated group (Fig. 9H and I). The results suggest that inhibiting PERK may mitigate TM-induced impairment of neuronal differentiation. Similarly, GSK2606414 significantly alleviated alcohol-induced impairment of neuronal differentiation, which was evident by increased formation of neurospheres and β-III-tubulin expression compared to EtOH-treated group (Fig. 10). On the other hand, GSK2606414 treatment failed to reverse TM-and alcohol-induced impairment of glial differentiation (Fig. 11).

**Figure 9.**
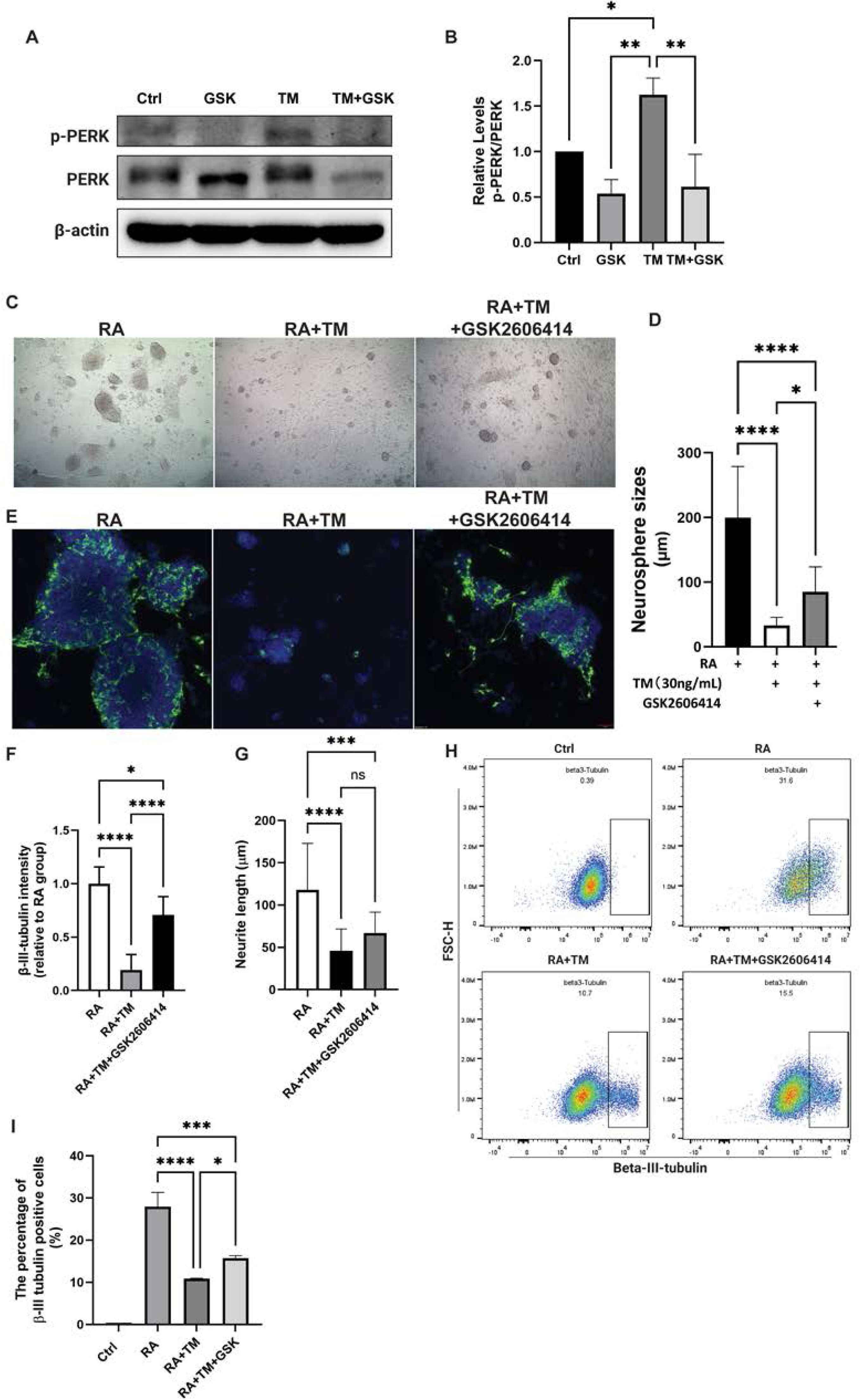
Effects of GSK2606414 on TM-induced impairment of neuronal differentiation. **A**: NE-4C cells were pretreated with 1 μM GSK2606414 or vehicle for 30 minutes, followed by the treatment with 30 ng/mL TM for 12 hours. Cell lysates were collected for IB analysis of p-PERK and PERK. **B**: The ratio of p-PERK/PERK was quantified. **C**: NE-4C cells were pretreated with 1 μM GSK2606414 for 30 min, followed by treatment with RA (1.5 × 10⁻L M) with or without 30 ng/mL TM for 48 hours. After 48 hours, the medium was replaced with RA-free medium. Cell morphology was examined under the bright-field imaging on day 3. **D**: The size of neurospheres was determined as described in Materials and Methods. **E**: The expression of β-III-tubulin in the cultures was determined by IF on day 3. **F**: The intensity of β-III-tubulin signals in panel **E** was quantified. **G**: The length of neurites was quantified. **H**: The β-III-tubulin positive cells were detected by flow cytometry on day 3. **I**: The percentage of β-III-tubulin-positive cells in panel **H** was quantified. Data are presented as mean ± SEM of three independent experiments. *p < 0.05, **p < 0.01. ***p < 0.001, **** < 0.0001 when compared to respective controls.

**Figure 10.**
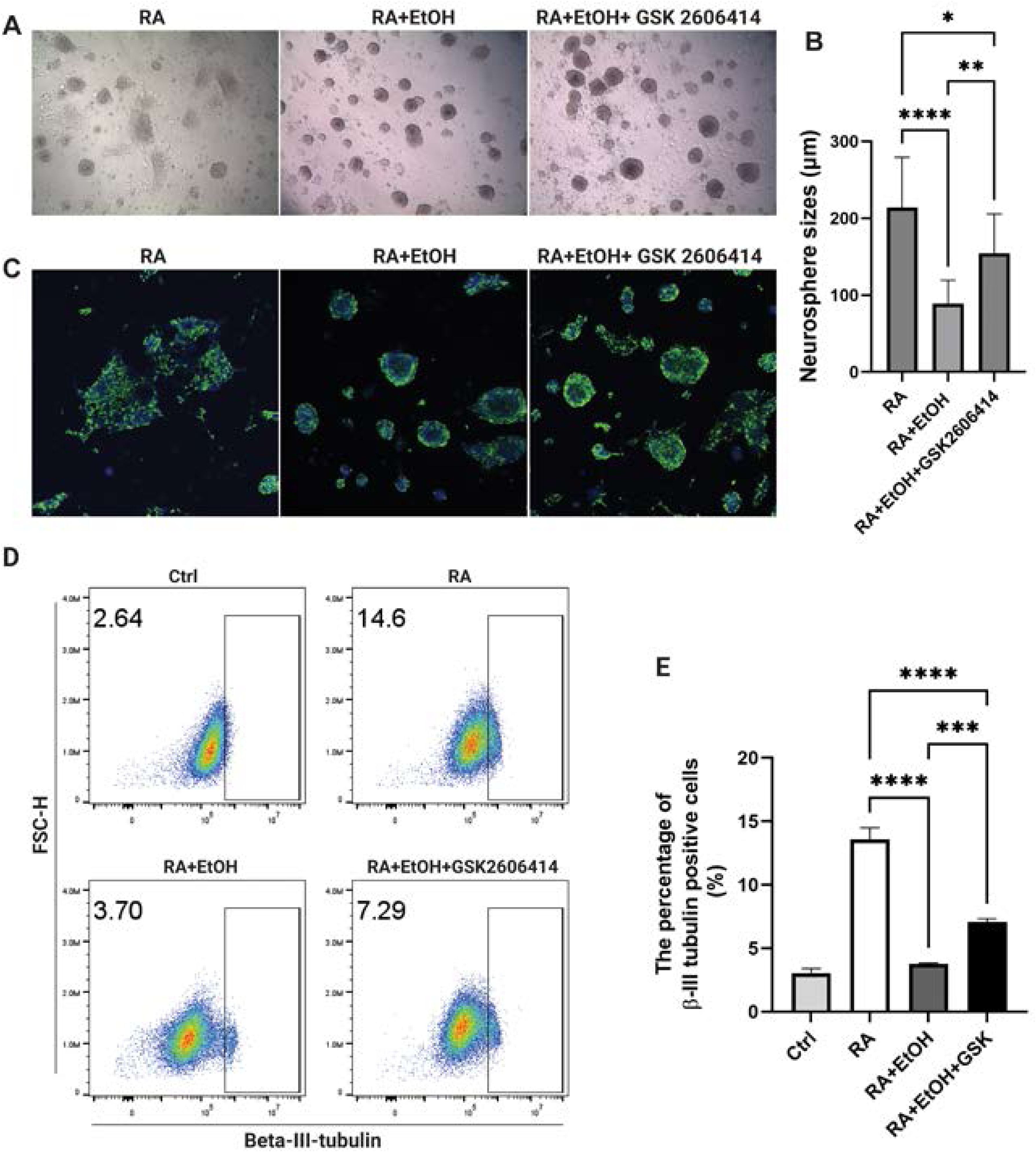
Effects of GSK2606414 on alcohol-induced impairment of neuronal differentiation. **A**: NE-4C cells were pretreated with 1 μM GSK2606414 for 30 min, followed by the treatment of RA (1.5 × 10⁻L M) with or without 0.4% alcohol (EtOH) for 48 hours. After that, the medium was replaced with RA-free medium and cell morphology was examined under the bright-field imaging on day 3. **B**: The size of neurospheres in panel **A** was quantified. **C**: The expression of β-III-tubulin on in the cultures was examined by IF on day 3. **D**: β-III-tubulin positive cells were determined by flow cytometry on day 3. **E**: The percentage of β-III-tubulin-positive cells in panel **D** was quantified. Data are presented as mean ± SEM of three independent experiments. *p < 0.05, **p < 0.01. ***p < 0.001, ****p < 0.0001 when compared to respective controls.

**Figure 11.**
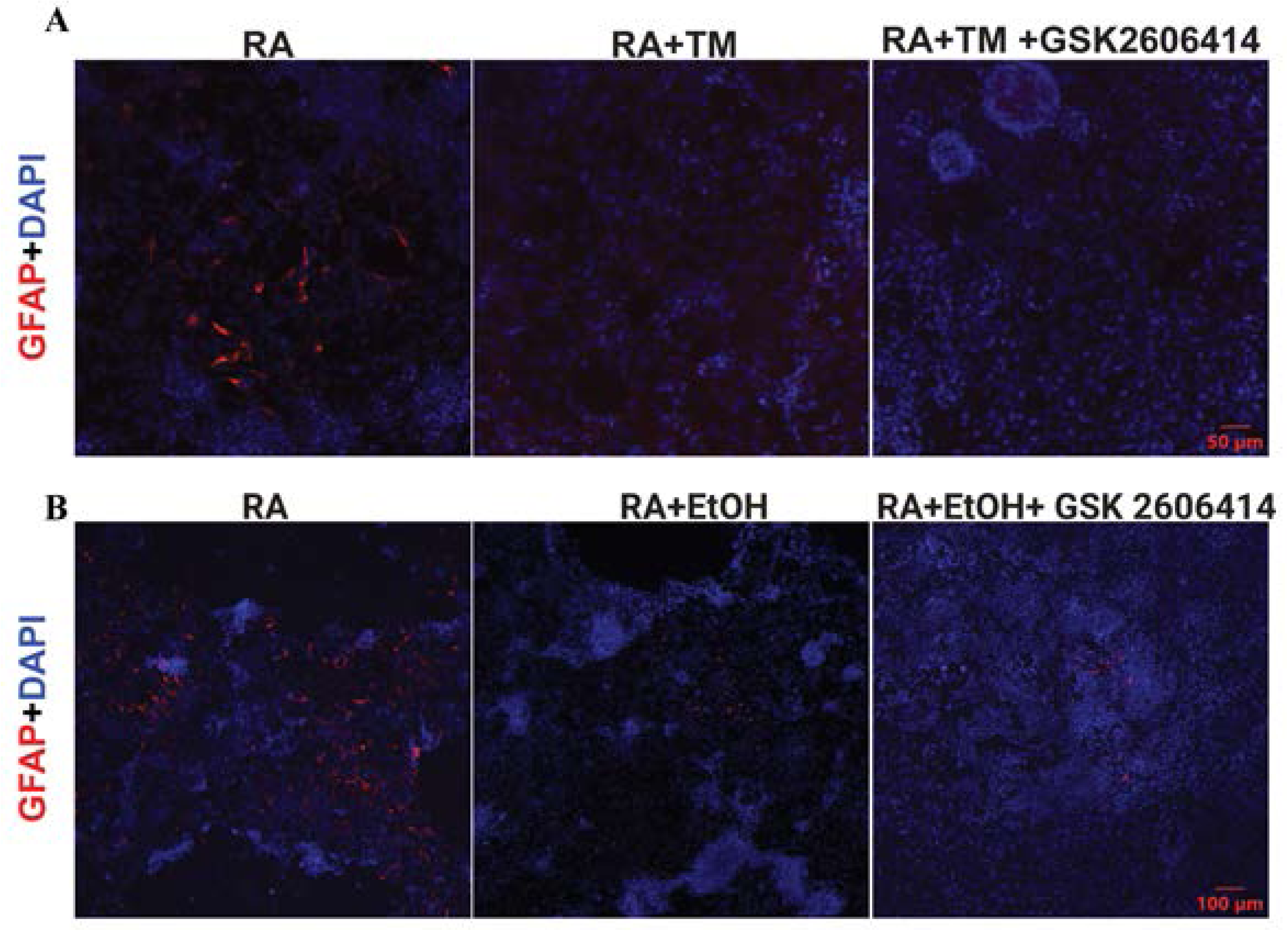
Effects of GSK2606414 on TM-and alcohol-induced impairment of glial differentiation. NE-4C cells were pretreated with 1 μM GSK2606414 for 30 min, followed by the treatment of RA (1.5 × 10⁻L M) with/without 30 ng/mL TM (**A**) or with/without 0.4% alcohol (EtOH)(**B**) for 48 hours. The medium was then replaced with RA-free medium. The expression of GFAP in the cultures was visualized by IF on day 9.

### Inhibiting PERK alleviates neuronal differentiation impairment caused by prenatal alcohol exposure (PAE) in mice

To further validate the *in vitro* findings and establish physiological relevance, we examined the effects of alcohol and GSK2606414 in pregnant mice. PAE has been commonly used to model FASD [34]. In our PAE paradigm, alcohol (5 g/kg) was administered to pregnant mice by oral gavage on GD14 and 16. The fetal brain tissues were collected on GD17. Consistent with *in vitro* data, PAE caused a significant upregulation of phosphorylated PERK in the GD 17 brain (Fig. 12 A and B). Alcohol did not affect the proliferative activity, which was indicated by comparable numbers of Ki67 positive cells in the VZ of alcohol exposed and control fetal brains (Fig. 12C and D). To determine whether PAE affect neuronal differentiation in the fetal brain and PERK activation is involved in this process, we administered GSK2606414 (50 mg/kg) to pregnant mice by oral gavage one hour prior to alcohol exposure, then examined the expression of phosphorylated PERK and apoptosis marker cleaved caspase-3 (Fig. 12E-G). GSK2606414 effectively inhibited PAE induced PERK activation in the fetal brain. Neither PAE nor GSK2606414 affect apoptotic cell death as indicated by little changes of cleaved caspase-3, suggesting that our PAE paradigm had minimal effects on cell survival in the fetal brain. We then assessed the effect of PAE and GSK2606414 treatment on neuronal differentiation by evaluating the number of cells expressing immature neuron marker doublecortin (DCX) in the ventricular zone (VZ). GSK2606414 partially but significantly reversed alcohol-induced reduction of DCX-positive cells (Fig. 12H and I). Since PAE and GSK2606414 had minimal effects on cell proliferation and survival, these data suggest that PAE may inhibit neuronal differentiation and GSK2606414 can partially reverse alcohol-induced inhibition. Taken together, these results were consistent with *in vitro* results and indicated that PERK played an important role in alcohol-induced impairment of neuronal differentiation.

**Figure 12.**
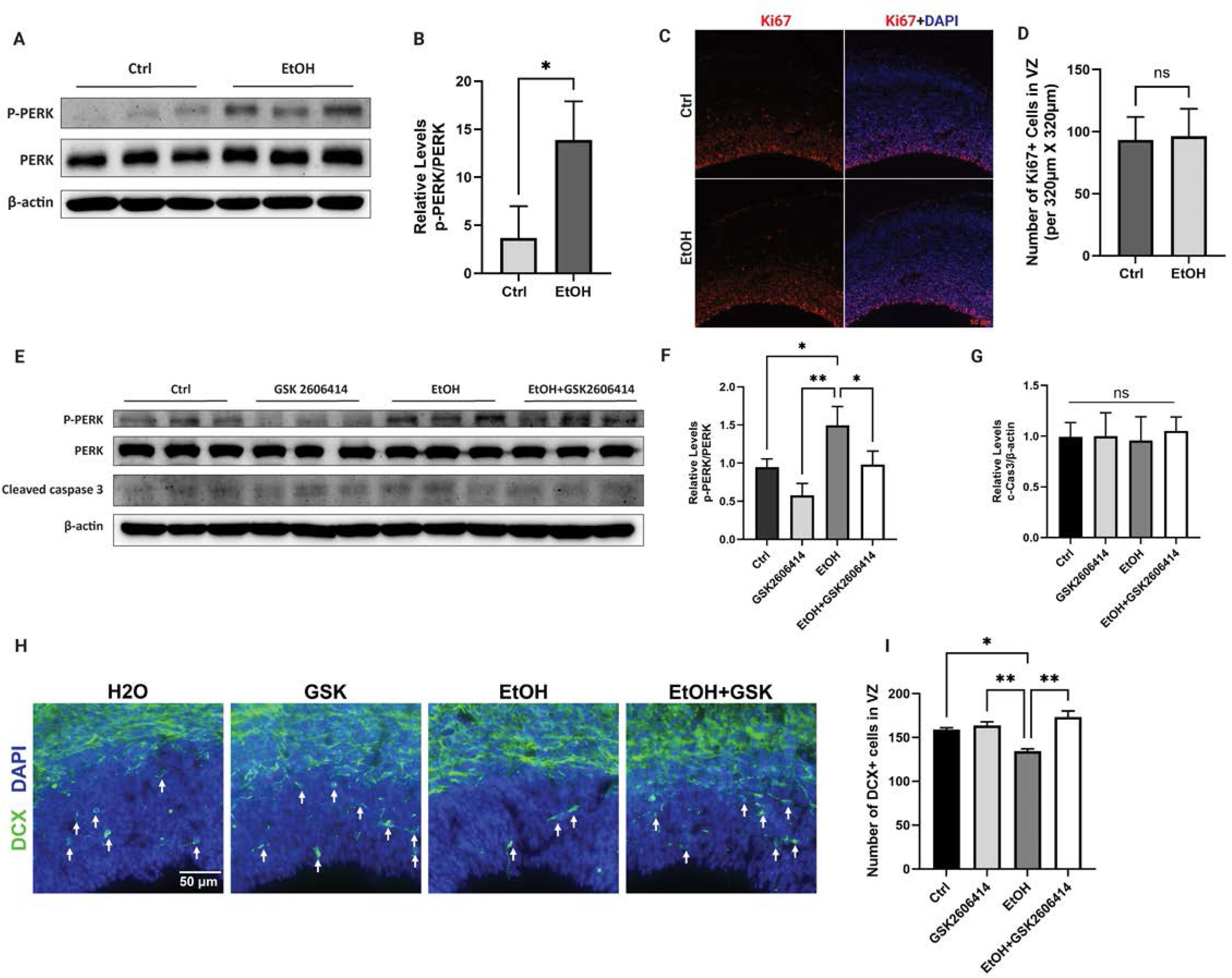
Effects of GSK2606414 on prenatal alcohol exposure (PAE)-induced impairment of neuronal differentiation in mice. **A**: Pregnant mice were either administered water or alcohol (EtOH: 5 g/kg) by oral gavage once per day on gestation days (GD) 14 and 16. On GD17, mice were sacrificed, and fetal brains were collected, and protein was extracted for IB analysis of p-PERK and PERK expression. **B**: The ratio of p-PERK/PERK was quantified. Data was expressed as mean ± SEM. n=3 per group. Data were analyzed by Student’s t test. *p< 0.05. **C**: GD17 fetal brain was sectioned for IF analysis of the expression of Ki67. **D**: The number of Ki67-positive cells was quantified. Data was expressed as mean ± SEM. n=3 per group. ns: not significant. **E**: Pregnant mice were pretreated with GSK2606414 (50 mg/kg) or vehicle 1 hour prior to the administration of water or alcohol (5 g/kg) by oral gavage once per day on GD14 and 16. On GD17, the mice were sacrificed, and fetal brains were collected. The protein was extracted and subjected to IB analysis of the expression of p-PERK, PERK, and cleaved caspase-3. **F**: The ratio p-PERK/PERK was quantified. Each data point was expressed as mean ± SEM. n=3 per group. The data were analyzed by One-way ANOVA followed by Tukey’s post hoc test. *p< 0.05; \*\**p*< 0.01.**G**: The levels of cleaved caspase-3 were quantified and normalized with β-actin. **H**: DCX-positive cells in the fetal brain were determined by IF on GD17. Arrows indicate the scattered DCX+ neurons in the ventricular zone (VZ). **I**: Quantification of the number of DCX+ neurons in the entire VZ from one coronal section per animal. Each data point was expressed as mean ± SEM. n=3 per group. Data were analyzed by One-way ANOVA followed by Tukey’s post hoc test. *p< 0.05; \*\**p*< 0.01.

## Discussion

In this study, we employed both *in vitro* and *in vivo* models to explore the mechanisms by which alcohol disrupts neural differentiation, a crucial process in neurogenesis and brain development. As an established *in vitro* model of neural differentiation, NE-4C cells—derived from the anterior brain vesicles of embryonic day 9 mouse embryos—represent proliferating neuroectodermal stem cells with the ability to differentiate into neurons and astrocytes upon RA induction. These cells are widely used to study the mechanisms of neural differentiation [16–18]. To model prenatal alcohol exposure (PAE) *in vivo*, we administered alcohol to pregnant mice via gavage at gestational days (GD) 14–16, a period equivalent to the human second trimester, which is a widely used experimental model for FASD [35–37]. Our findings revealed that alcohol impaired the RA-induced differentiation of NE-4C cells into neurons and astrocytes without affecting cell migration. Additionally, alcohol exposure induced ER stress, specifically activating the PERK pathway. In the PAE model, the number of DCX+ cells-a marker of newly differentiated immature neuron—was reduced by alcohol exposure, while cell death and proliferation were largely unaffected. Notably, inhibiting PERK rescued alcohol-induced deficits in neuronal differentiation in NE-4C cells and mitigated the reduction of DCX+ cells in the VZ of fetal brain following PAE. Collectively, these results suggest that alcohol disrupts neural differentiation by triggering ER stress, particularly through activation of the PERK pathway.

Our findings demonstrating alcohol-induced impairment of neuronal differentiation align with previous reports showing that alcohol disrupts neurogenesis by inhibiting neuronal differentiation and altering neuronal morphology [38–43]. This consistency supports the relevance of our model system for investigating the cellular and molecular mechanisms underlying these effects. Similarly, our results indicating that alcohol inhibits astrocyte differentiation agree with prior research [44–47]. For instance, Vallés et al. [47] reported that PAE delayed GFAP expression in the rat brain and radial glial cell cultures. Additional studies have shown that in utero alcohol exposure significantly reduces GFAP expression in both the embryonic cerebral cortex [45] and postnatal brain [46]. However, in models of postnatal alcohol exposure, alcohol appears to increase GFAP expression in rodents and human brain organoids [48–50], suggesting that its effects may vary depending on the timing of alcohol exposure [51]. In our model, alcohol did not impact neural stem cell (NSC) migration, whereas some studies suggest that alcohol may impair NSC and neural progenitor cell motility by disrupting cytoskeletal organization [52–54]. These discrepancies highlight the importance of considering key variables such as the timing of alcohol exposure, model systems, and alcohol dosage when evaluating its effects on CNS development.

Alcohol is reported to induce ER stress in various cell types of different organ systems, which is implicated in the pathogenesis of multiple diseases associated with alcohol exposure [5–8]. For example, in the liver, alcohol exposure induced ER stress which playing a crucial role in the progression of alcoholic liver disease (ALD)[55]. Alcohol drastically upregulated UPR proteins which ultimately lead to hepatocyte injury and liver fibrosis [7]. Similarly, alcohol-induced ER stress is implicated in pancreatic injury and alcoholic pancreatitis, where excessive UPR activation contributed to pancreatic inflammation and dysfunction of pancreatic cells [6]. It is also well-established that alcohol-induced ER stress in cardiomyocytes resulted in cell death and contributed to alcoholic cardiomyopathy [5, 56].

ER stress has been implicated in the development of various neurodegenerative diseases and neurodevelopmental disorders [57–62]. Our previous research demonstrated that excessive ER stress induced by alcohol exposure can contribute to neurodegeneration, suggesting a role for ER stress in alcohol-induced neurotoxicity [11, 12]. However, whether ER stress is involved in alcohol’s disruption of neural stem cell (NSC) differentiation remains unclear. Our current findings support the hypothesis that alcohol impairs neural differentiation through ER stress, based on several key observations. First, alcohol-induced disruption of neural differentiation was accompanied by ER stress activation, particularly the PERK pathway. Second, treatment with TM, an ER stress inducer that inhibits N-glycosylation and leads to the accumulation of misfolded proteins, similarly impaired neural differentiation. Third, MANF deficiency in NE-4C cells which activated ER stress and the PERK pathway also impaired neural differentiation. As a well-known ER-resident protein, MANF plays a crucial role in maintaining ER homeostasis, and its deficiency has been shown to induce ER stress in neuronal cells. Finally, inhibiting PERK, a key component of the unfolded protein response (UPR), significantly mitigated alcohol-induced impairment of neural differentiation.

Given the high demand for protein synthesis during NSC differentiation, maintaining proper proteostasis is essential for this process. Compelling evidence suggests that ER stress and the unfolded protein response (UPR) play a crucial role in NSC differentiation. For instance, a previous study reported that ER stress disrupted neuronal differentiation and inhibited neurite outgrowth in mouse embryonic carcinoma P19 cells [63]. Additionally, UPR activation has been shown to impair the balance between direct and indirect neurogenesis, leading to premature neuron generation [64]. Interestingly, some UPR proteins are upregulated during the neuronal differentiation of mouse embryonic stem cells [65], suggesting that the increased demand for protein synthesis during NSC differentiation may itself trigger ER stress. Alternatively, specific UPR proteins may actively participate in regulating neural differentiation.

In response to ER stress, the UPR is activated through three key branches: PERK, IRE1, and ATF6. The specific activation of these pathways depends on the type of stressor, the cellular context, and the intensity of stress. Initially, the UPR functions to restore ER homeostasis by reducing global protein synthesis, enhancing protein folding capacity, and facilitating the degradation of misfolded proteins. However, when ER stress is excessive or prolonged, UPR signaling shifts toward pro-apoptotic pathways, ultimately leading to cell dysfunction and death. As discussed earlier, UPR may also influence NSC differentiation. In our model, alcohol preferentially activated the PERK/eIF2α pathway in NSCs. Similarly, ER stress induced by TM or MANF deficiency strongly stimulated the PERK/eIF2α pathway, further supporting its role in alcohol-induced impairment of neural differentiation.

Unlike the ATF6 and IRE1 branches of the unfolded protein response (UPR), which are primarily protective under ER stress, the PERK branch plays a dual role—it can either restore cellular homeostasis or initiate cell death pathways. The PERK-eIF2α pathway is critical for regulating protein synthesis and cellular stress adaptation. Upon activation, PERK phosphorylates eIF2α at Ser51, leading to a global reduction in protein translation to alleviate ER burden while selectively enhancing the translation of ATF4. ATF4 orchestrates a transcriptional response that promotes amino acid metabolism, oxidative stress resistance, and autophagy [66–69]. However, under prolonged or severe ER stress, ATF4 induces CHOP (C/EBP Homologous Protein), which triggers apoptotic cell death [70–72]. This pathway also includes a feedback mechanism via GADD34, which dephosphorylates eIF2α to restore normal protein synthesis once the stress is resolved. The PERK/eIF2α pathway is thus a key determinant of cell fate under ER stress conditions.

The involvement of the PERK/eIF2α pathway in neurogenesis has been previously documented. For instance, during corticogenesis, activation of PERK/eIF2α has been linked to microcephaly and a reduction in neural progenitor populations in mice [64]. To investigate its role in alcohol-induced impairment of neural differentiation, we employed PERK inhibition. As most well-characterized PERK inhibitor, GSK2606414 is potent and pharmacokinetically suitable for both *in vitro* and *in vivo* studies [73–76]. Our results showed that GSK2606414 effectively inhibited alcohol-induced PERK activation *in vitro* and *in vivo*. More importantly, inhibiting PERK activity significantly mitigated alcohol-induced impairment of neuronal differentiation, supporting the involvement of the PERK/eIF2α pathway in this process.

In our PAE model, which corresponds to the human second trimester, we observed that PAE inhibits neuronal differentiation, as evidenced by a reduction in DCX-positive immature neurons within the ventricular zone (VZ) of the fetal brain. DCX (doublecortin) is a microtubule-associated protein expressed in migrating immature neurons, and its expression is restricted to cells committed to the neuronal lineage [77]. In the embryonic mouse brain, DCX is prominently expressed in immature neurons migrating within the VZ [78]. Although β-III-tubulin is widely used as a neuronal marker due to its expression in postmitotic neurons [22, 23], it is not optimal for assessing neurogenesis at embryonic day 17 (E17) in mice. By this developmental stage, β-III-tubulin is widely and intensely expressed, making it difficult to distinguish newly generated neurons. Therefore, DCX was employed as a more suitable marker for identifying newly formed neurons in this study. Our results demonstrate that PAE significantly reduces the population of DCX-positive immature neurons in the VZ. Notably, these effects appear to involve the PERK/eIF2α signaling pathway. Future studies should investigate whether pharmacological inhibition of PERK can mitigate PAE-induced neurobehavioral deficits in offspring. Such work would not only further validate the involvement of the PERK/eIF2α pathway but also assess its potential as a therapeutic target for preventing alcohol-induced neurodevelopmental impairments.

Notably, inhibition of PERK activity did not reverse the impairment of glial differentiation induced by either alcohol or TM, suggesting that the PERK/eIF2α pathway is not directly involved in this aspect of neurodevelopmental disruption. However, findings from our ER stress models—including alcohol exposure, TM treatment, and MANF deficiency—consistently indicate that ER stress adversely affects glial differentiation. To further evaluate the role of ER stress in alcohol-induced impairment of glial development, future studies may employ broader ER stress inhibitors, such as the FDA-approved chemical chaperone 4-phenylbutyric acid (4-PBA). Previously, we utilized 4-PBA to investigate the contribution of ER stress to alcohol-induced neurotoxicity in the postnatal mouse brain, supporting its potential utility in delineating ER stress-mediated mechanisms in glial lineage specification [79].

The precise mechanism by which the PERK/eIF2α pathway regulates NSC differentiation remains to be fully elucidated. One plausible mechanism involves the pathway’s role in translational control: activation of PERK leads to phosphorylation of eIF2α, resulting in global attenuation of protein synthesis—a process critical for proper neural differentiation. In addition to PERK, three other kinases—protein kinase R (PKR), heme-regulated inhibitor kinase (HRI), and general control nonderepressible 2 kinase (GCN2)—can also phosphorylate eIF2α. Together, these kinases constitute the integrated stress response (ISR). It would be of interest to investigate whether activation of these alternative ISR pathways similarly disrupts NSC differentiation. The ISR inhibitor ISRIB, which selectively reverses the effects of eIF2α phosphorylation, represents a promising tool for probing this question. Notably, ISRIB has been shown to ameliorate spatial learning and memory deficits in rodent models following adolescent alcohol exposure, further supporting its potential relevance in mitigating alcohol-induced neurodevelopmental impairments [80].

## Abbreviation

AD: Alzheimer’s disease
ASD: autism spectrum disorder
ATF4: activating transcription factor
ATF6: activating transcription factor
CHOP: C/EBP homologous protein
CNS: central nervous system
DCX: doublecortin
eIF2α: eukaryotic initiation factor 2α
ER: endoplasmic reticulum
EtOH: ethanol
FASD: fetal alcohol spectrum disorder
GD: gestation day
GFAP: glial fibrillary acidic protein
GRP78: glucose-regulated protein 78
IRE1α: inositol-requiring enzyme 1α
MANF: mesencephalic astrocyte-derived neurotrophic factor
NSCs: neural stem cells
PAE: prenatal alcohol exposure
PD: Parkinson’s disease
PERK: PKR-like ER kinase
RA: retinoic acid
ROS: reactive oxygen species
TM: tunicamycin
UPR: unfolded protein response
VZ: ventricular zone

## Acknowledgement

This work was supported by the National Institutes of Health (NIH) grants AA017226 and AA015407. We thank Jianyu Yu for his assistance in the experiment of flow cytometry analysis.

---

**Supplementary Figure 1**. Effects of tunicamycin on the viability of NE-4C cells. NE-4C cells were treated with different concentrations of tunicamycin (TM) for 72 hours. MTT assay was performed to determine the cell viability. The experiments were replicated three times (n = 3). Each data point is presented as mean ± SEM. **p < 0.01, when compared to control.

**Supplementary Figure 2**. Effects of PERK inhibitor GSK2606414 on the viability of NE-4C cells. NE-4C cells were treated with different concentrations of GSK2606414 for 72 hours. Cell viability was assessed using the MTT assay. Each data point is presented as mean ± SEM of three replications. ***p < 0.001, when compared to control.

