## Supplementary figures and images for "Alcohol Disrupts Neural Differentiation Through Endoplasmic Reticulum Stress and PERK Pathway Activation"

### Supplementary Figure 1

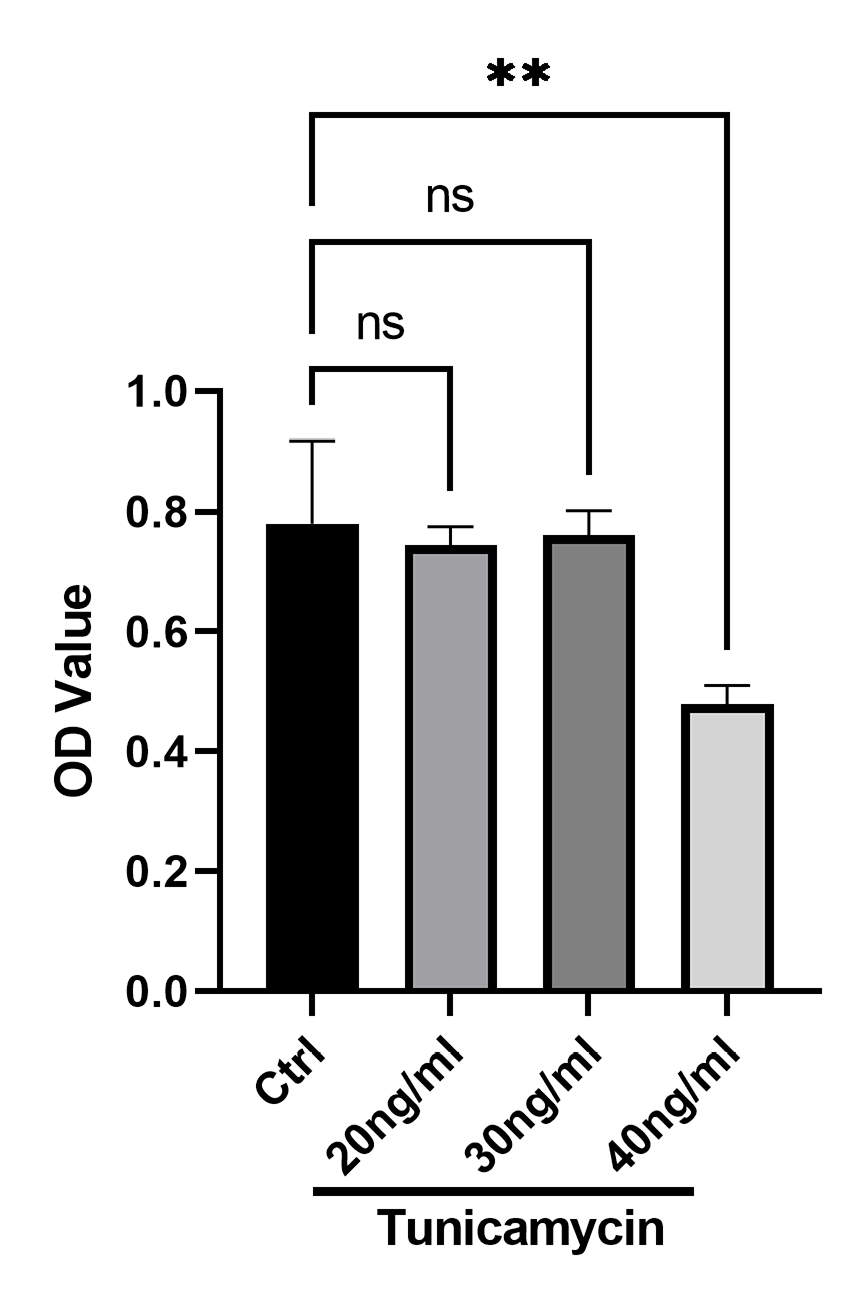

### Supplementary Figure 2

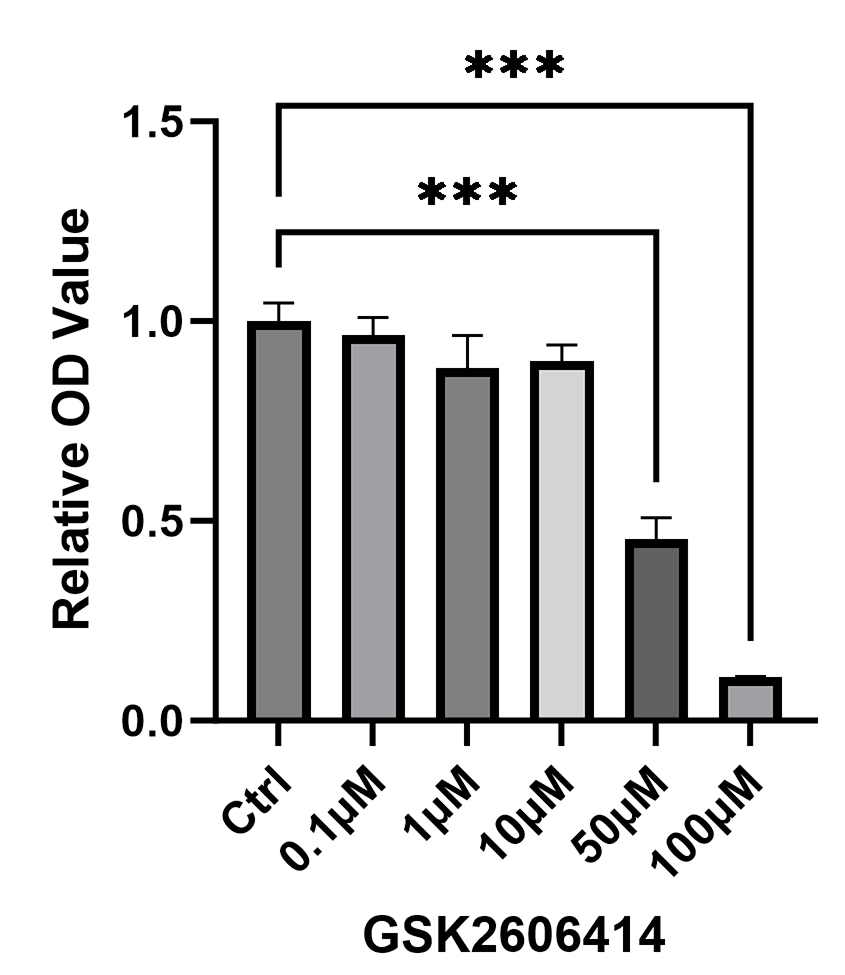
